# Discovery of peptides for targeted delivery of mRNA lipid nanoparticles to cystic fibrosis lung epithelia

**DOI:** 10.1101/2023.09.13.557559

**Authors:** Melissa R. Soto, Mae M. Lewis, Jasmim Leal, Yuting Pan, Rashmi P. Mohanty, Sophie Peng, Tony Dong, Debadyuti Ghosh

## Abstract

For cystic fibrosis (CF) patients, a lung targeted gene therapy would significantly alleviate pulmonary complications associated with morbidity and mortality. However, mucus in the airways and cell entry pose huge delivery barriers for local gene therapy. Here, we used phage display technology to select for and identify mucus- and cell-penetrating peptides against primary human bronchial epithelial cells (pHBECs) from CF patients cultured at air-liquid interface (ALI). At ALI, pHBECs produce mucus and reflect CF disease pathology, making it a clinically relevant model. Using this model, we discovered a lead candidate peptide, and incorporated it into lipid nanoparticles (LNPs) to deliver mRNA to pHBECs and mouse lungs *in vivo*. Compared to LNPs without our peptide, peptide-LNPs demonstrated 7.8-fold and 4.8-fold higher mRNA expression *in vitro* and *in vivo*, respectively. Since gene delivery to pHBECs is a significant challenge, we are encouraged by these results and anticipate that our peptide could be used to successfully deliver CF gene therapies in future work.

## Introduction

Cystic Fibrosis (CF) is a genetic disease caused by single mutations within the cystic fibrosis transmembrane conductance regulator (CFTR) gene, and as a result, requires gene therapy to correct the mutation and restore CFTR function. While these mutations affect all organs, mucus-build up found in the lungs of CF patients results in the most severe lung morbidities which are the leading cause of death for CF patients.(1) As a result, many efforts focus on developing gene therapy formulations that can be locally delivered to the lungs.(2) Local administration to the pulmonary tract is attractive since it allows for higher, safer dosing than feasible with systemic delivery.(3) However, the hostile environment found in CF patients’ lungs, which includes hyperconcentrated mucus and immune cells such as macrophages, greatly hinders treatments from reaching intended target sites (i.e., cells harboring the CFTR mutation). Also, negatively charged therapeutic nucleic acids cannot enter target cells on their own, which makes intracellular delivery a challenge. Given that the thick mucus layer and cell membrane are two of the biggest barriers to successful gene delivery, it is necessary to develop gene therapy carriers that can overcome these obstacles.

Traditionally, clinics have used virus vectors for CF gene therapy due to their intrinsic ability to enter mammalian cells. However, their use can be limited due to their innate immunogenicity, broad natural tropism for nonspecific cells and tissues, and limited size capacity for packaging nucleic acids.(2) Due to these shortcomings, non-viral delivery systems are alternative carriers but are often road-blocked by the various extracellular and intracellular barriers of uptake (i.e., mucosal and cell membrane). To overcome the mucus barrier, a common approach researchers use to enhance non-viral delivery systems is to formulate with mucus-penetrating polymers, such as polyethylene glycol (PEG). However, PEGylation can hinder cellular uptake and PEG alone does not have inherent cell-targeting capabilities.(4–6) From these considerations, peptides are a promising alternative to polymers for gene therapy formulations. Peptides are attractive delivery systems since they can shuttle therapeutics to their intended target site, can potentially enhance cell internalization, and be easily incorporated into non-viral delivery systems where larger payloads can be used.(7) Based on their amino acid sequence, peptides can possess a wide range of desirable properties, to facilitate mucus penetration, and importantly, they have been used to demonstrate cell-specific targeting and uptake, which address the two major barriers mentioned before.

To achieve targeted delivery, researchers have used directed evolution to select virus variants with redirected tropism or discover cell-targeting peptides through technologies such as phage display. For example, groups have evolved adeno-associated viruses and lentiviruses to target human CF epithelia in order to improve the transduction of mammalian viral delivery systems.(8–10) For phage display, on the other hand, M13 or T7 bacterial viruses, i.e., phage, are traditionally used as screening tools to discover peptides with desired properties. Here, phage are genetically engineered to display recombinant random peptides or proteins on their viral coat proteins (i.e., capsid proteins). For instance, T7 phage can be genetically modified to display up to 415 copies of a single peptide on the surface of their capsids and phage libraries can be created that consist of up to 10^9^-10^10^ unique peptide-presenting phage or “clones”. Pooled together, these random peptide-presenting phage libraries can be incubated or “panned” against a target of interest. Through iterative rounds of panning against the target, there is a selection pressure applied such that peptide(s) with the desired properties are identified. After identification, peptides are typically removed from the structural context of the phage and incorporated into delivery systems (e.g., antibody, lipid nanoparticles, viruses) to redirect targeting.(11) In the context of respiratory targeting, multiple groups have demonstrated the utility of phage display for discovering peptides that can target the lungs, both systemically and locally.(12–15) For CF therapy specifically, Romanczuk et al, panned phage display libraries against primary human bronchial epithelial cells (pHBECs), however, panning was done with pHBECs from non-CF donors in solution, meaning these cells were not fully demonstrative of CF airway conditions *in vivo*.(12)

For CF gene therapy, it is necessary to identify ligands that overcome the mucus and the cellular barrier to facilitate targeted delivery to the affected CF airway epithelia. While in earlier work we identified mucus-penetrating peptides by panning our custom T7 phage display library against an *in vitro* CF-like mucus model,(16) this strategy focused on mucus penetration in a synthetic disease model of CF and did not fully address the challenges of targeting and entering the airway epithelia in clinically relevant CF disease models. To address these critical issues, we leveraged directed evolution with phage display to discover and select for peptides with the desired physicochemical properties to achieve both mucus and cell penetration. Here, we used T7 phage display to pan against fully differentiated pHBECs isolated from CF patients, which produce mucus and have ciliated cells, and phenotypically and physiologically represent a more disease relevant model to mimic conditions found in the lungs of CF patients.(17) With this strategy, our goals were to identify peptides and their physicochemical properties and confirm they improve phage uptake into mucus-producing pHBECs. Additionally, we aimed to demonstrate that select peptide candidates could be incorporated into clinically relevant lipid nanoparticles (LNPs) to improve mRNA delivery and expression *in vitro* in fully differentiated CF patient cells and *in vivo* in lungs via pulmonary delivery.

## Results

### Selection of enriched mucus and cell-penetrating T7 phage clones

For our selection strategy, we iteratively panned a cysteine constrained T7 7-mer peptide-presenting phage library against mucus producing primary human bronchial epithelial cells (pHBECs). Specifically, we used pHBECs cultured in physiologically relevant air-liquid interface (ALI) to identify phage that can overcome two main delivery barriers in the airway space for CF therapy: 1) mucosal barrier and 2) cellular barrier (Fig. 1). We chose pHBECs isolated from CF patients cultured on ALI as our selection model since it more accurately reflects the CF microenvironment upon differentiation (i.e., mucus production and subsequent ciliary movement). An overview of our selection strategy is depicted in Figure 1A. To increase selection pressure throughout our rounds of panning, we sequentially shortened incubation times from rounds 1 to 3 and increased the time cells were incubated in wash buffer from round 1 to round 2 (selection details from each round are shown in Figure 1B). We next quantified internalized phage after each round of selection by standard double-layer plaque assay to determine enrichment following each round of selection. After five rounds of selection, we saw an average of 380-fold increase in C_out/Cin_ (Concentration of phage internalized or “output”/Concentration of input phage), demonstrating that our CX7C T7 library was enriched.

**Figure 1.**
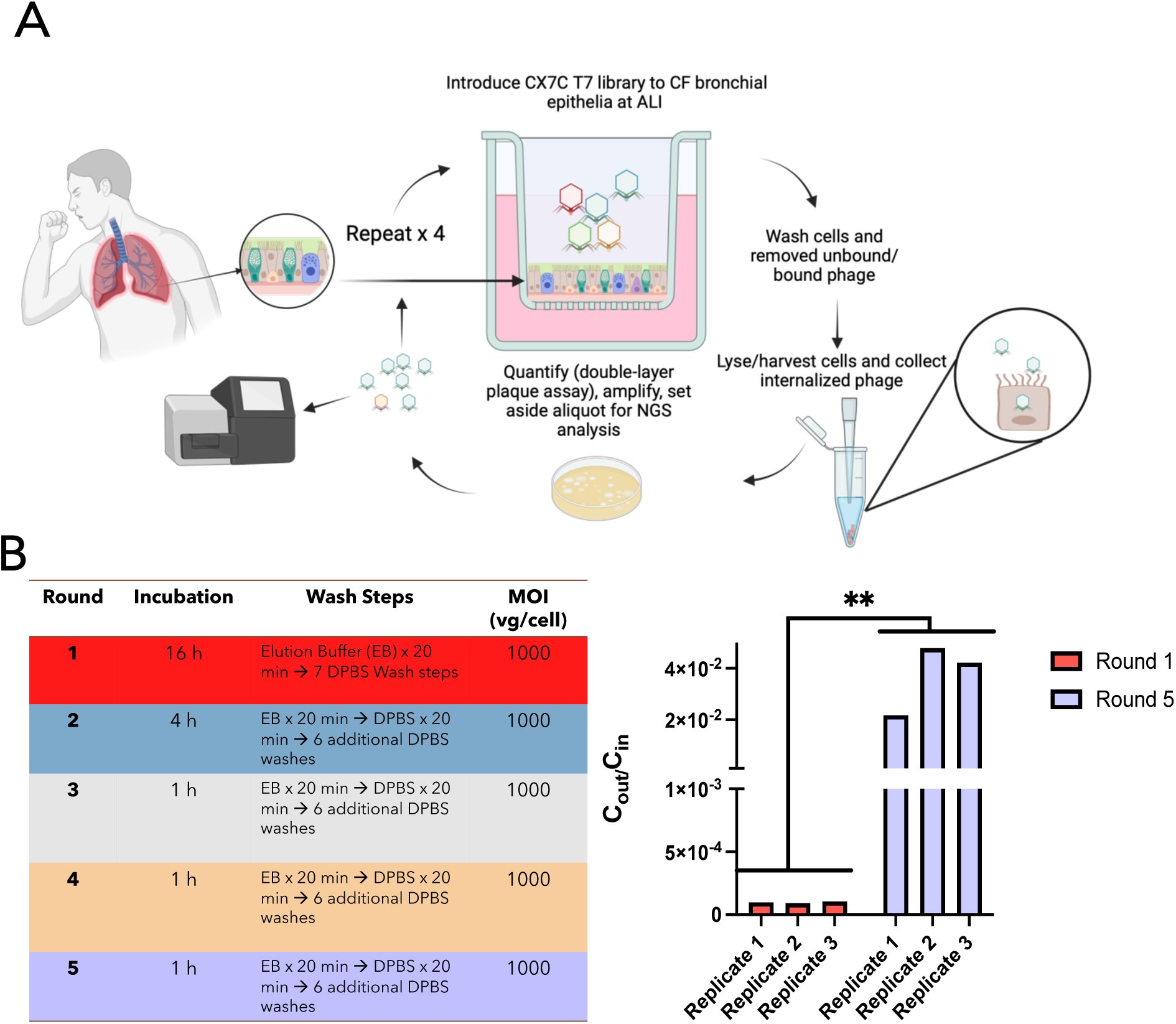
Panning strategy and enrichment. **(A)** Selection strategy using a cysteine constrained random 7-amino acid peptide T7 phage display library (CX7C) against differentiated primary human bronchial epithelial cells (pHBECs) through iterative, high-throughput selection for a total of five rounds. Image made with BioRender. **(B)** Selection conditions (left) and enrichment (right) of CX7C phage library for enhanced CF pHBEC uptake. C_out_ represents the concentration of phage collected after each round and C_in_ is the initial input phage concentration. Phage were added at 1000 viral genomes/cell (vg/cell) and incubated for 16 hours at 37°C for round 1 and 1 hour at 37°C for round 5. Data represents change in C_out_/C_in_ for three separate replicates from the first to last round of selection. Unpaired t-test *\*\*p < 0.01*.

### Enriched clones display peptides that are positively charged and hydrophilic

Using Next Generation Sequencing (NGS) and a custom python script (16, 18) for data analysis, we identified “hits” or the top 30 most abundant peptide sequences from each round of selection and ranked these peptides from 1 to 30 *(N=*3). For each ranked peptide *(N=*3), we calculated and compared its net charge in rounds 1 and 5 to determine if there was a shift in net charge after all rounds of selection. A heat map depicting these values indicates that the net charge of peptides shifted to a slightly more positive value from round 1 to round 5 (Figure 2A).

**Figure 2.**
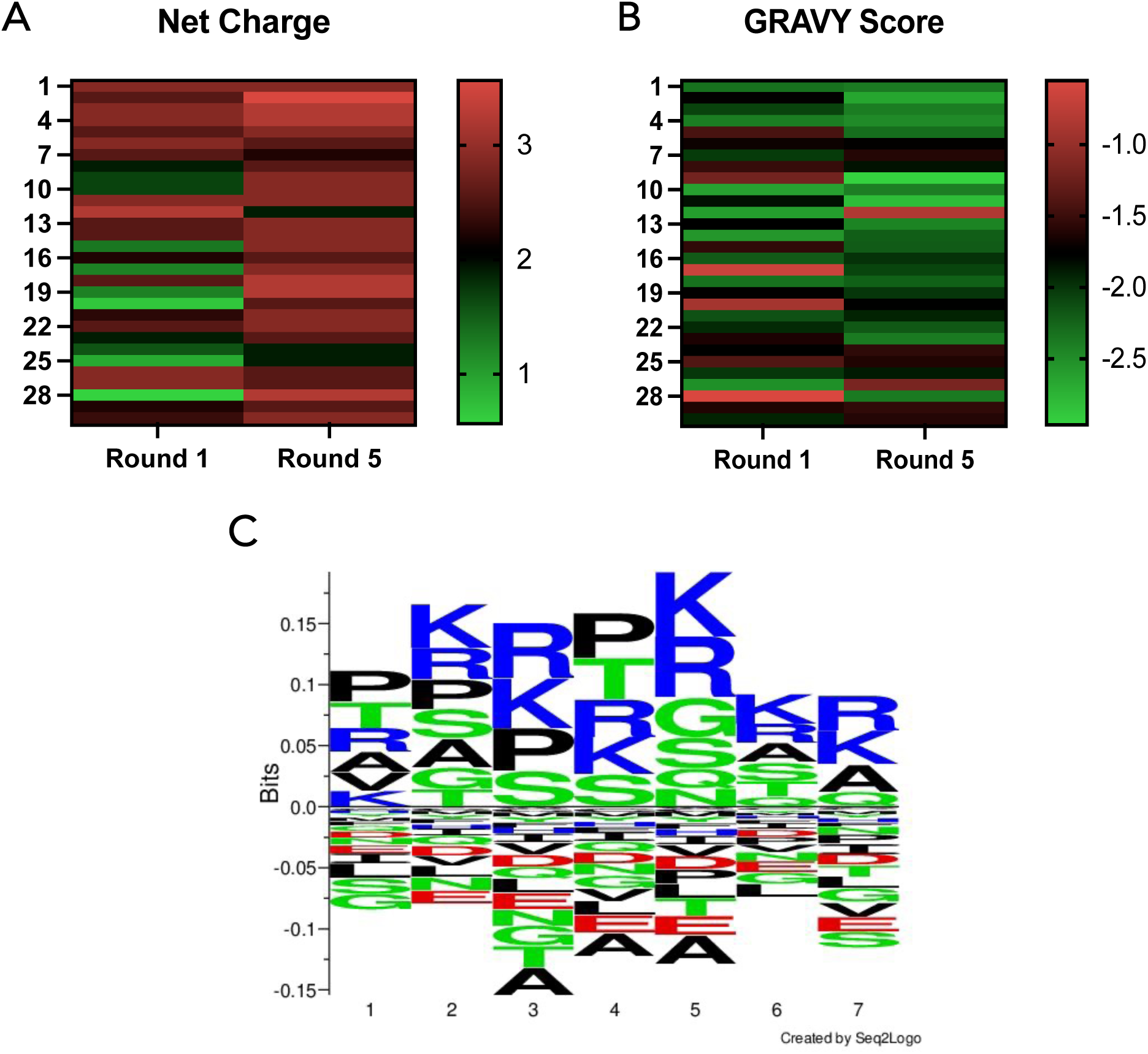
Physicochemical properties of peptides displayed on enriched phages. **(A)** Net charge and **(B)** GRAVY score values were calculated for the top 30 most frequent peptide sequences from each replicate *(N=*3); total of 90 sequences analyzed per round). Peptides were ranked 1-30 based on frequency. Graphed are the mean values for each ranked peptide sequence *(N=*3). **(C)** Multiple sequence alignment visual representation using Seq2Logo for the top 30 sequences of each replicate after 5 rounds of selection.(19)

Similarly, we calculated and compared the grand average of hydropathicity, or GRAVY score, from the top 30 most frequently occurring peptide sequences from rounds 1 and 5 (Fig. 2B). The more positive the GRAVY score, the more hydrophobic the peptide sequence, and vice versa. Here, we also observed a shift from the first to last round of selection to a slightly more negative GRAVY score, indicating that there were more hydrophilic sequences present after 5 rounds of selection (Fig. 2B).

Additionally, we implemented the Seq2Logo tool to visualize a multiple sequence alignment for the top 30 sequences of all replicates *(N=*3) after the 5^th^ round and identify the amino acid frequency at each position of the 7-mer peptide.(19) We observed that this consensus sequence was rich in the basic and hydrophilic amino acids arginine and lysine, which further supported trends we saw for net charge and GRAVY scores (Fig. 2C).

Lastly, we further analyzed these sequences to select individual clones for further validation. From each replicate, we selected clones that were within the top 10 most abundant sequences in the 5^th^ round and found in all three replicates (Table 1). We observed that each clone showed an increase in their abundance, i.e., percentage of total sequences identified from NGS analysis after each subsequent round (Fig. 3). The enrichment of relative frequency of individual clones ranged from ∼7.7 (Clone B) to 13.7 (Clone C) from the first round to fifth round. Except for Clone F, net charges were 2.9 and GRAVY scores were less than −2.0. Clone F had a net charge and GRAVY score of 0.9 and −0.26, respectively.

**Table 1.**
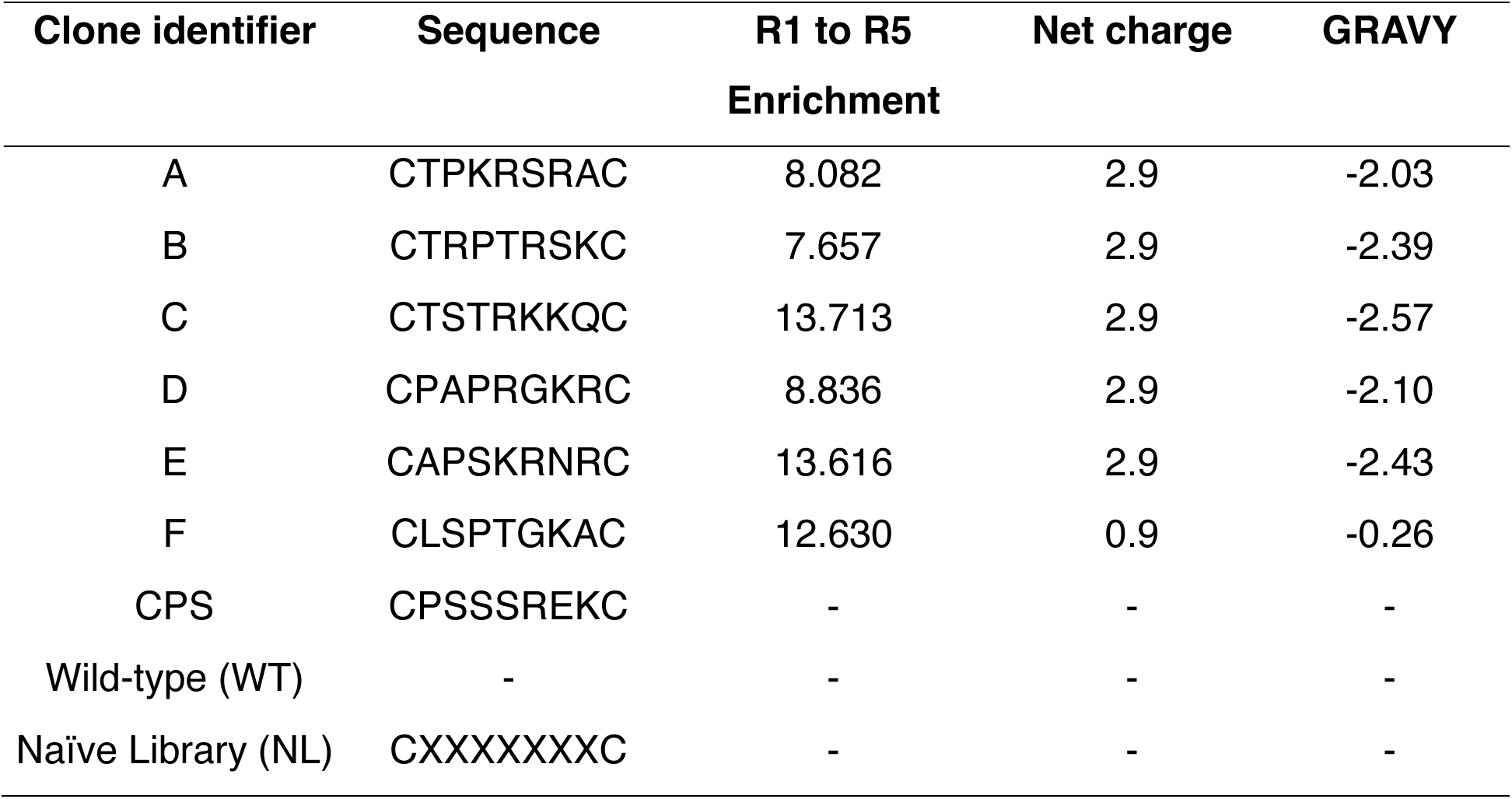
T7 phage clones selected for validation and the physicochemical properties of their displayed peptides.

**Figure 3.**
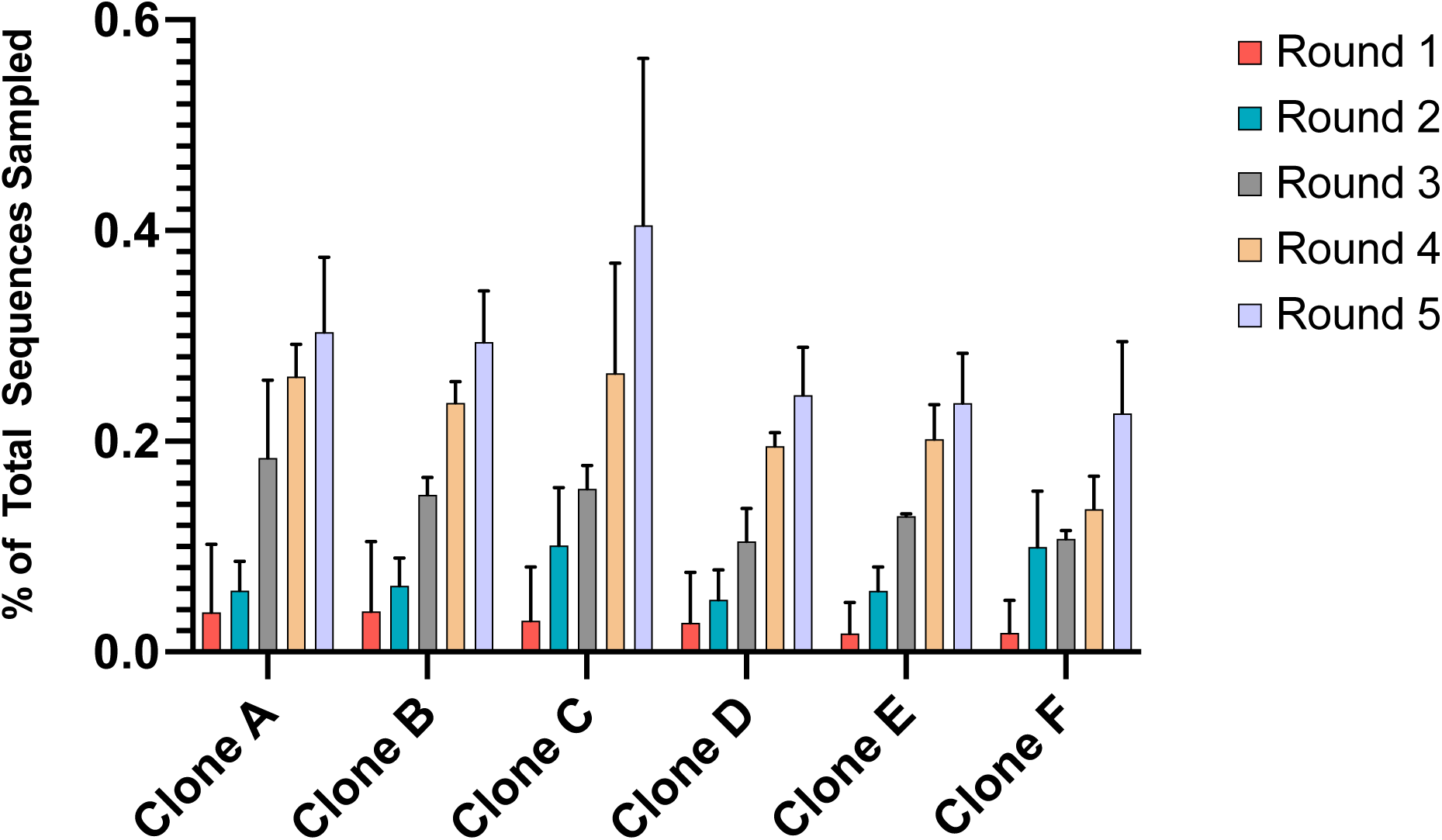
Enrichment of individual clones identified using high throughput sequencing data obtained from NGS. We identified peptide-displaying clones that were present in the top 10 most abundant sequences across all three replicates. For each round of panning, the % of total sequences sampled represents the (average peptide frequency count/total # of sequences obtained from NGS data) x 100.

### Enriched clones demonstrate improved transport through mucus producing CF pHBECs

Like our panning procedure, we differentiated our pooled pHBECs (see Materials and Methods) and incubated them with the individual peptide-presenting phage clones that we identified during the selection process (Table 1) to confirm that these clones achieve both mucus penetration and cell uptake (Fig. 4). From our validation study, we found that of the six clones validated, clones B-E demonstrated significant difference in uptake compared to an unselected naïve library control (*p<0.05)*. Additionally, Clones B, C, and E had significantly improved uptake compared to a peptide insertless “WT” control and an internal control sequence (denoted as “CPS”) (Table 1). Clone CPS was previously discovered in our lab from phage display screening for improved diffusion through CF-like mucus.(16) Our best performing clone, Clone C, demonstrated a ∼454-fold (*p<0.0005)*, 56-fold (*p<0.005)*, and 50-fold (*p<0.005)* improvement in uptake compared to Naïve library, WT, and Clone CPS.

**Figure 4.**
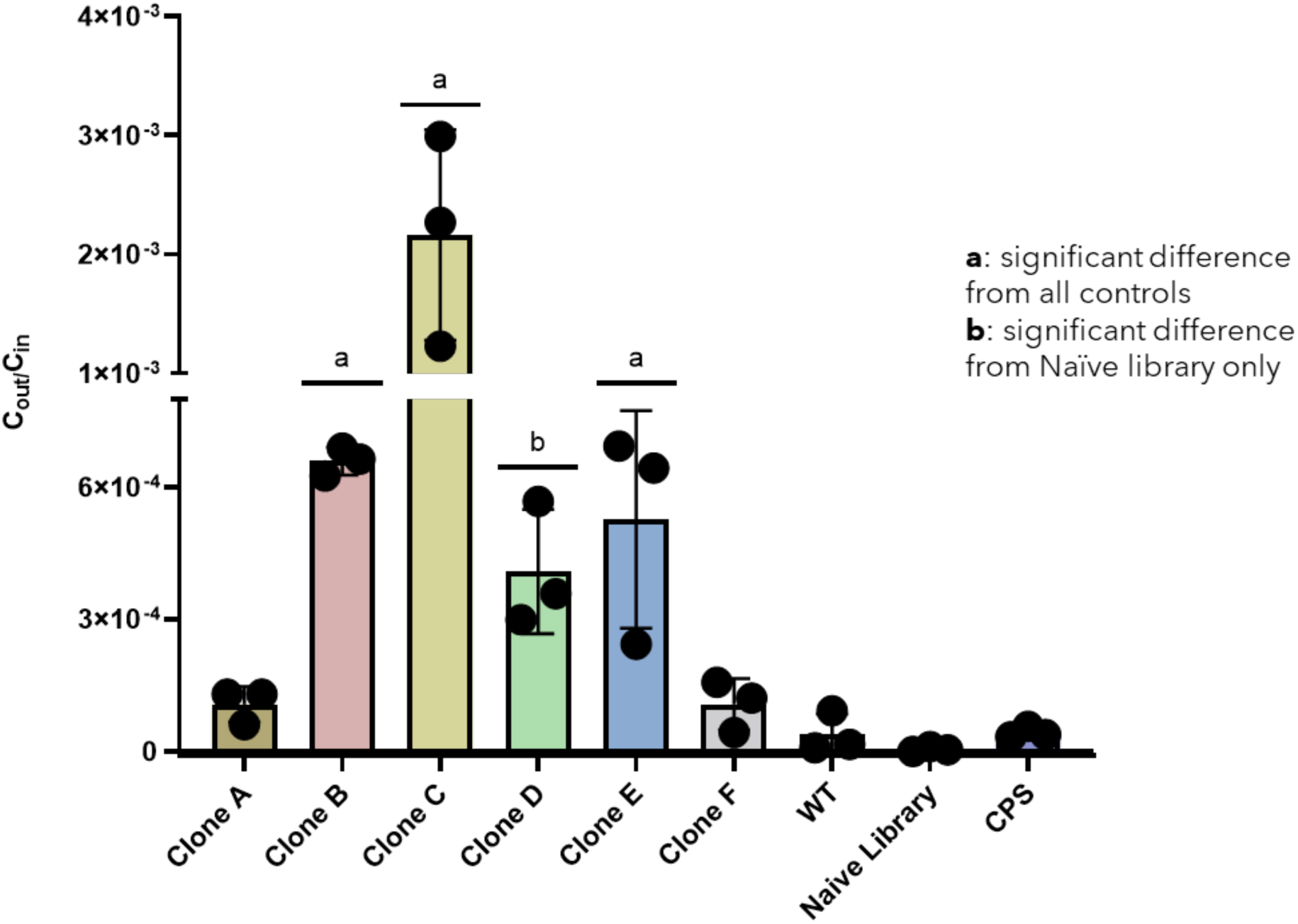
Validation of selected clones from panning. C_out_ represents the output concentration of phage collected from ALI cells after 1h and C_in_ is the initial input phage concentration. Controls include wild-type (WT) phage which lacks peptides on its capsid surface, Naïve library which is a mixture of 7-mer peptide presenting phage clones, and a positive mucus-penetrating clone displaying the peptide CPSSSREKC (net charge 1 and GRAVY score of −1.2). One-way ANOVA (Kruskal-Wallis test; uncorrected Dunn’s); *a,b p<0.05*.

### Peptides can be successfully incorporated into lipid nanoparticles (LNPs)

We confirmed that peptides could improve transport and intracellular uptake in CF pHBECs while in the context of a phage particle. Next, we sought to determine whether these peptides could be effectively incorporated into a lipid nanoparticle (LNP) formulation. We encapsulated nanoluciferase (NLuc) reporter mRNA into LNPs (NLuc LNPs) via microfluidic mixing using the four lipids used in Moderna’s FDA-approved Spikevax formulation (Table 2). Based on its abundance and validation in phage uptake, we selected Peptide C from Clone C to introduce into LNPs for subsequent studies. We used either Peptide C or a control mucus-penetrating peptide, CPS, conjugated to a myristic acid fatty acid as peptide-lipids to incorporate into LNPs as a fifth component lipid. Peptide-lipids or additional PEG (to serve as an additional mucus-penetrating control) were added to Moderna’s Spikevax base composition at an optimized percentage (Table 2).

**Table 2.**
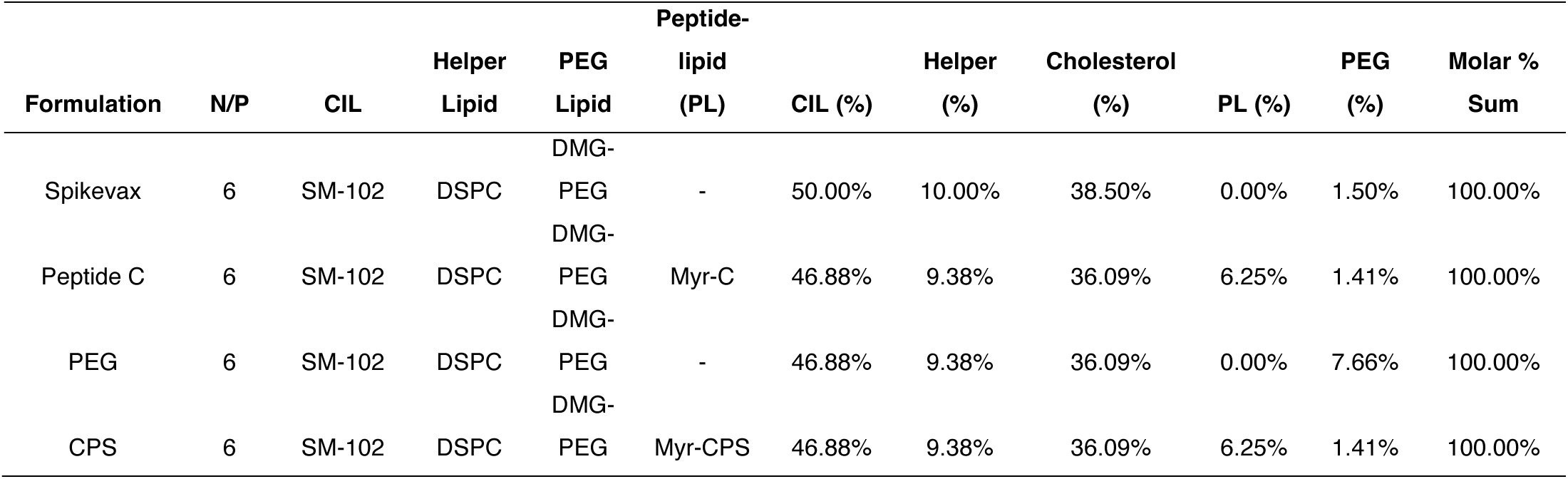
LNP formulation details. N/P = nitrogen to phosphate ratio; CIL = cationic ionizable lipid; PEG = polyethylene glycol; DMG-PEG = 1,2-dimyristoyl-rac-glycero-3-methoxypolyethylene glycol-2000; DSPC = Distearoylphosphatidylcholine; Myr = myristoyl group

After formulating LNPs, dynamic light scattering data showed that all formulations had diameter sizes below 80 nanometers. The polydispersity index (PDI), which approximates the monodispersity of the formulations, was highest for our PEG formulation, while all other formulations had PDIs < 0.2. Next, we characterized their encapsulation efficiencies (EEs); all but the PEG LNP formulation resulted in EEs > 90%.

### Selected peptide functionalized LNPs enhance mRNA expression

After confirming that peptides could be incorporated into LNPs, we determined how each formulation affected mRNA expression. We delivered NLuc-LNPs to differentiated pHBECs and incubated for 48h before measuring bioluminescence. Here, we saw significantly enhanced mRNA expression between Peptide C-LNPs and all other formulations (Fig. 6). Notably, our Peptide C-LNP demonstrated 7.8-fold higher expression than Moderna’s Spikevax LNP.

To determine if our Peptide C-LNPs demonstrated preferential uptake in primary HBECs, we measured mRNA expression in an alternative cell model, a THP-1 derived macrophage cell line. Using a similar incubation time (i.e., 48 hours) to our validation study on pHBECs, we observed significantly lower mRNA expression (1.7-fold) from Peptide C-LNP groups compared to Spikevax LNP treatment groups. Expression was slightly lower than our CPS-LNP group and slightly higher (but not statistically different) than the PEG treated group (Supplementary Fig. 1).

### Selected peptide functionalized LNPs enhance mRNA expression in vivo

Next, we sought to determine how our Peptide C could enhance delivery and mRNA expression *in vivo*. We delivered LNP formulations intratracheally, and after 24 hours, harvested and separated each lung into their five separate lobes (i.e., left lung, right cranial lobe, right accessory lobe, right caudal lobe, and right middle lobe). We observed a similar trend to our *in vitro* ALI uptake study – our Peptide-C formulation group showed the highest bioluminescence, followed by CPS-LNP, Spikevax-LNP and lastly PBS group (Figure 7). Compared to Spikevax, Peptide C-LNPs demonstrated 4.4-fold higher bioluminescence. Notably, we also observed mRNA expression distributed throughout all lobes for our Peptide C-LNP group (Figure 7).

## Discussion

For diseases affecting the lungs, such as cystic fibrosis (CF), local delivery of therapeutics is promising since it results in fewer systemic side effects, requires a lower dose compared to systemic delivery, and therapeutics can be administered in multiple ways that are amenable to outpatient use (e.g., nebulizer, dry powder, metered-dose inhaler).(3, 20–23) For example, when comparing inhalation versus oral delivery of the same drug substance, lung: plasma drug concentration ratios can reach over 100 (compared to 1.6 for oral administration), making local delivery advantageous for targeting the lung tissues.(23) For CF, this pharmacokinetic profile is attractive for the delivery of gene therapies like CFTR gene replacement or correcting specific CFTR mutations. However, both mucus and cell barriers hinder transport and delivery of therapeutics into the cell.(24–29) To improve intracellular uptake, phage display can be useful for selecting targeting ligands with affinity for cells and specific tissues. We previously used T7 phage combinatorial libraries against a CF-like mucus model and screened for peptides that showed enhanced transport through CF patient-derived sputum.(16) While this initial work demonstrated that peptides could be used as coatings to improve transport through mucus models, there remained critical questions to address. With this novel screening strategy, while we successfully identified mucus-penetrating peptides, we did not pan for ligands that can both facilitate mucus penetration and also achieve the necessary targeted uptake into the CF-affected airway epithelial cells; both barriers are critical bottlenecks to achieve nucleic acid delivery needed for gene therapy. Also, our previous screening was against CF-like mucus with components to best mimic CF clinical samples; however, recent advances have shown that primary cells isolated from CF patients and grown under appropriate conditions (i.e., air-liquid interface (ALI)) are more accurate physiological models of the disease. As a result, we built upon our previous work and used primary cells derived from CF patients as our selection model. Here, we used fully differentiated primary cells homozygous for the deltaF508 mutation to better reflect this disease state in the human lung. At ALI, these clinically obtained cells can produce mucus, ciliate, and even mimic CF pathology more closely than traditionally used cell lines. (30–36)

To ensure that candidate ligands facilitated intracellular uptake, we modified our panning strategy to collect peptide-presenting phage internalized by fully differentiated CF epithelia (Fig. 1). This approach is in contrast to others’ phage display work that primarily focused on collecting and isolating cell surface bound phage on either undifferentiated primary HBE cells or HBE cell lines.(12, 14, 37) With our collection method, we increased the probability that only internalized peptide-presenting phage were enriched prior to subsequent rounds in the selection process (Fig. 1). Compared to our earlier work,(16) we also greatly modified our panning to increase selection pressure at various rounds (Fig. 1B); during selection, the increase in stringency (either by reducing incubation time and/or increasing washes) should allow selection of the ‘winners’ that can survive in each replicate. In other words, we improved the probability of discovering peptides with increased propensity to get into cells.(38) This enrichment of peptide-presenting phage is indicative of affinity-based selection, as indicated by other groups.(8, 38, 39) After five rounds of selection, we observed 380-fold enrichment (Fig. 1) which aligns with our previous studies that utilized similar T7 phage libraries.(16, 18)

Next, we sought to characterize the physicochemical properties of these enriched peptides. We found that sequences in the fifth round were more positively charged and hydrophilic than those in the first round (Fig. 2A). This shift in hydrophilicity was reflected in our previously published work, and supports other findings that suggest hydrophilicity aids in mucus diffusion.(16, 40) Additionally, this finding corroborates extensive work by others showing that poly(ethylene) glycol (PEG) polymer is an attractive coating to improve mucus penetration of carriers due to its hydrophilicity.(6, 41–47) For example, Schneider et al. showed that PEGylated nanoparticles have higher mucus diffusion in CF sputum *ex vivo* and more uniform distribution in mouse airway epithelia compare to non-PEGylated nanoparticles. We did observe a change in the net charge with each subsequent round of selection with more positively charged peptide sequences (Fig. 2B). Previously, when we screened against CF-like mucus only, we saw peptide sequences that were more negative to neutral net-charged.(16) One potential and likely explanation for this observed difference is that our enriched peptides mimicked more cell-penetrating peptides, which are positively charged in nature, in order to bind and be internalized by cells.(7, 48, 49) Importantly, we observed that our sequences do not possess a high enough positive charge such that the binding would be too strong with the net negative charged mucus microenvironment and be stuck for hindered transport.(16, 50) For instance, our recent work demonstrated that peptides with a net charge of 5 (at pH 7) have hindered diffusion through CF sputum.(16) This finding supported other work that has been shown in other extracellular net negative charge environments such as cartilage extracellular matrix that a high enough positive charge peptide would be immobilized. Given that the net charge of our lead peptide candidate is around 2.9, it is probable that the charge is positive enough to allow for weak partitioning through the mucus and eventually achieve cellular uptake.(50, 51) However, others have shown that net charge alone is not sufficient to determine transport and that spatial configuration and sequence order of amino acids also contributes to transport through the mucus barrier.(50) Future studies are warranted to investigate how sequence specificity affects mucus penetration and cellular uptake.

For validation studies, this strategy allowed us to proceed with six peptide sequences that were present in the top 10 most abundant peptides and present in all three replicates from our panning (Table 1). For each of these selected sequences, there was an increased affinity for the cells as observed by an increased frequency of the individual clones present in each subsequent round of selection (Fig.3). This enrichment indicated to us that there was an ‘evolutionary’ pressure to select, and not just screen, for peptides with desired properties for mucus penetration and cell uptake.(38) Additionally, after each round, this improvement in our selection process was validated where all peptide-presenting phage clones selected for validation performed better, i.e., achieved greater cellular uptake, compared to a peptide-less control, the original, unselected naïve library, and previously discovered mucus-penetrating peptide presenting phage (Fig. 4). Here, the only difference between all phage clones was the peptide sequence (or lack of) present on the phage surface. Given this detail, we expect these differences in uptake are based solely on capsid surface properties, which is supported by our other related work.(16, 18)

After demonstrating the efficacy of our peptide displayed on the phage capsid, we ultimately wanted to use our targeting peptide ligands to aid gene delivery across mucus and cell barriers. Therefore, we also confirmed that our leading peptide candidate demonstrated similar efficacy when introduced to a synthetic carrier. Here, we incorporated our peptide (conjugated to a fatty acid tail) into a clinically relevant lipid nanoparticle (LNP) system as an additional fifth component into Moderna’s Spikevax COVID-19 mRNA vaccine base formulation loaded with NLuc mRNA (Table 2). Cheng et al. have demonstrated that a fifth component lipid can be used to supplement LNPs (which typically consist of four lipid components) and interestingly, depending on the physicochemical property of the lipid, facilitate targeting to specific organs (i.e., lung, liver, spleen).(52) Additionally, others have incorporated peptides in nanoparticle or nanocomplex formulations in order to improve transfection or gene delivery to certain cell types.(47, 53–58) Notably, others have demonstrated that tandem peptides (i.e., peptides attached to lipid tails) containing targeting peptides can enhance transfection to neuronal and cancerous cell lines.(53–55) In these studies, tandem peptides were complexed or self-assembled with both RNA and protein cargos. For our work, we decided to leverage these advancements by integrating peptides conjugated to myristic acid (14-carbon fatty acid) into LNPs with standard microfluidic mixing.(54, 55) Here, we confirmed that our peptide C-lipid could be co-formulated with a conventional 4-component LNP and result in acceptable size (diameter < 100 nm), are monodisperse (polydispersity index < 0.2), and high encapsulation efficiency (>95%) (Fig. 5).(59) To validate whether incorporation of our peptide into LNPs could improve nucleic acid delivery to enhance protein expression, we evaluated LNP uptake into cells in differentiated pHBECs and measured luminescence using the Nano-Glo^®^ Luciferase assay system. After a 48-hour incubation period, we observed 10.5-fold and 4.5-fold increased luminescence from mRNA delivered by our Peptide C-LNP compared to LNPs containing additional PEG or a CPS-conjugated lipid, respectively, which were included as mucus-penetrating LNP controls (Fig. 6A). These results supported others’ findings that the addition of polymers used only for mucus-penetration may not be sufficient for also overcoming the cellular barrier.(6) For example, Conte et al., tested and compared PEGylated and non-PEGylated nanoparticles as siRNA carriers for CF treatment. They found that while PEGylated nanoparticles had improved transport through an artificial mucus model, they resulted in lower cellular uptake in a Calu-3 ALI cell model.(6)

**Figure 5.**
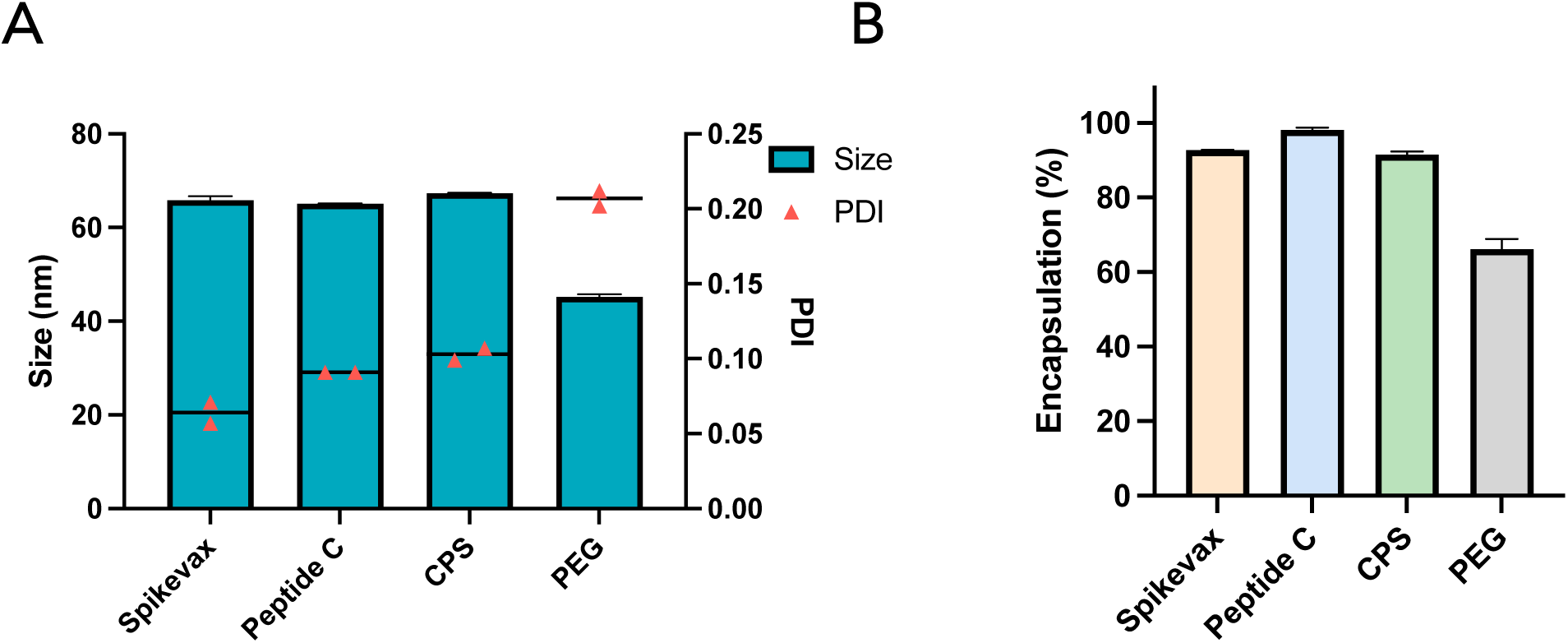
Characterization of LNPs. We determined size (in nm) and polydispersity index (PDI) by Dynamic Light Scattering (DLS) *(N=*2) **(A)** and encapsulation efficiencies **(B)** by modified Ribogreen assay *(N=*2).

**Figure 6.**
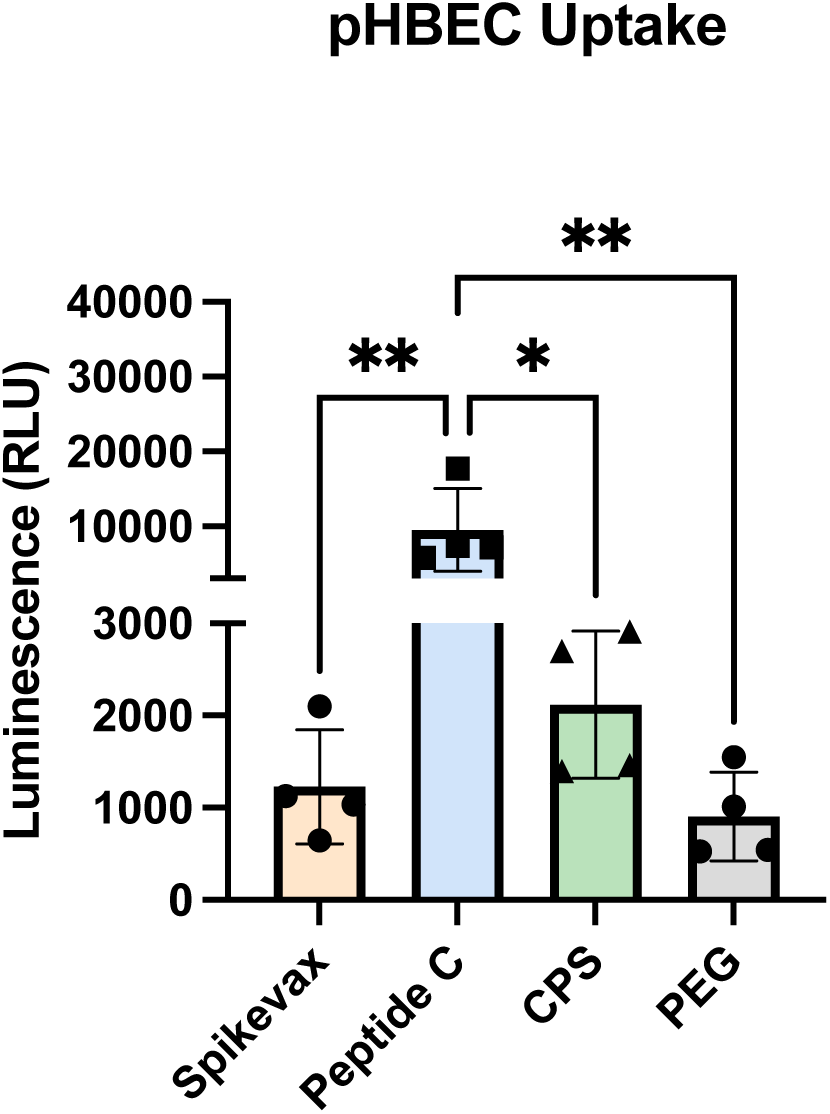
NLuc mRNA expression in primary HBECs following LNP treatment. We added LNPs (450 ng of mRNA) to the apical side of differentiated primary HBECs (*N*=4 per treatment group) and incubated them at 37°C for 48 hours. Following incubation, we measured bioluminescence by plate reader. One-way ANOVA (Tukey’s multiple comparisons test); *\* p<0.05, ** p <0.01*.

Nucleic acid delivery to the lungs is not only limited by mucus and cell barriers, but also immune cells present in the airways, especially in diseased lungs.(60, 61) Immune cells such as macrophages are present for immunosurveillance during homeostasis, and in CF sputum they can be present in larger numbers compared to a healthy patient.(62, 63) Given immune cell presence in the lungs, we sought to determine if our peptides were selective towards targeted uptake to CF patient epithelia by repeating the same Nano-Glo^®^ assay with THP-1 derived human macrophage cell line. In THP-1 cells, we observed lower luciferase expression from Peptide C-LNP relative to Spikevax and CPS-LNP formulations (Supplementary Fig. 1). Considering that Peptide C-LNP achieved substantially higher mRNA delivery than other LNPs in pHBECs from CF patients, these results suggest that Peptide C has enhanced uptake in pHBECs, relative to the Spikevax “peptide-less” control, compared to macrophages. This result is significant since inhaled nanoparticles, (e.g., gold nanoparticles) can be cleared by macrophages that reside in the lung.(64–67)

While we have demonstrated that our peptide-LNPs can achieve targeted mRNA delivery in a relevant *in vitro* airway epithelia model of CF, we next wanted to investigate delivery in preclinical models that can better provide the dynamics, anatomical structure and physiology of the lungs. Here, we delivered LNPs intratracheally to the airways of Balb/c mice to determine if our leading peptide candidate could enhance mRNA expression *in vivo*. Following 24 h after delivery of LNPs encapsulating NLuc mRNA, we observed significantly higher luminescence compared to all other groups (Fig. 7). Interestingly, this trend paralleled what we observed *in vitro* using differentiated pHBECs, where luminescence was highest for Peptide C-LNP followed by CPS-LNP and lastly Spikevax LNP (Fig. 6). Multiple groups have recently used NLuc (and modified NLuc) mRNA to optimize LNP formulations, even for respiratory delivery.(68–73) For instance, Lokugamage et al., optimized LNPs for nebulization using a modified NLuc reporter gene and demonstrated that their lead LNP could successfully deliver therapeutic mRNA as well. This highlighted that it is possible to replace NLuc reporter mRNA with a more clinically relevant cargo and still produce significant results. This also encourages us to exchange our NLuc cargo with therapeutic mRNA in future work.

**Figure 7.**
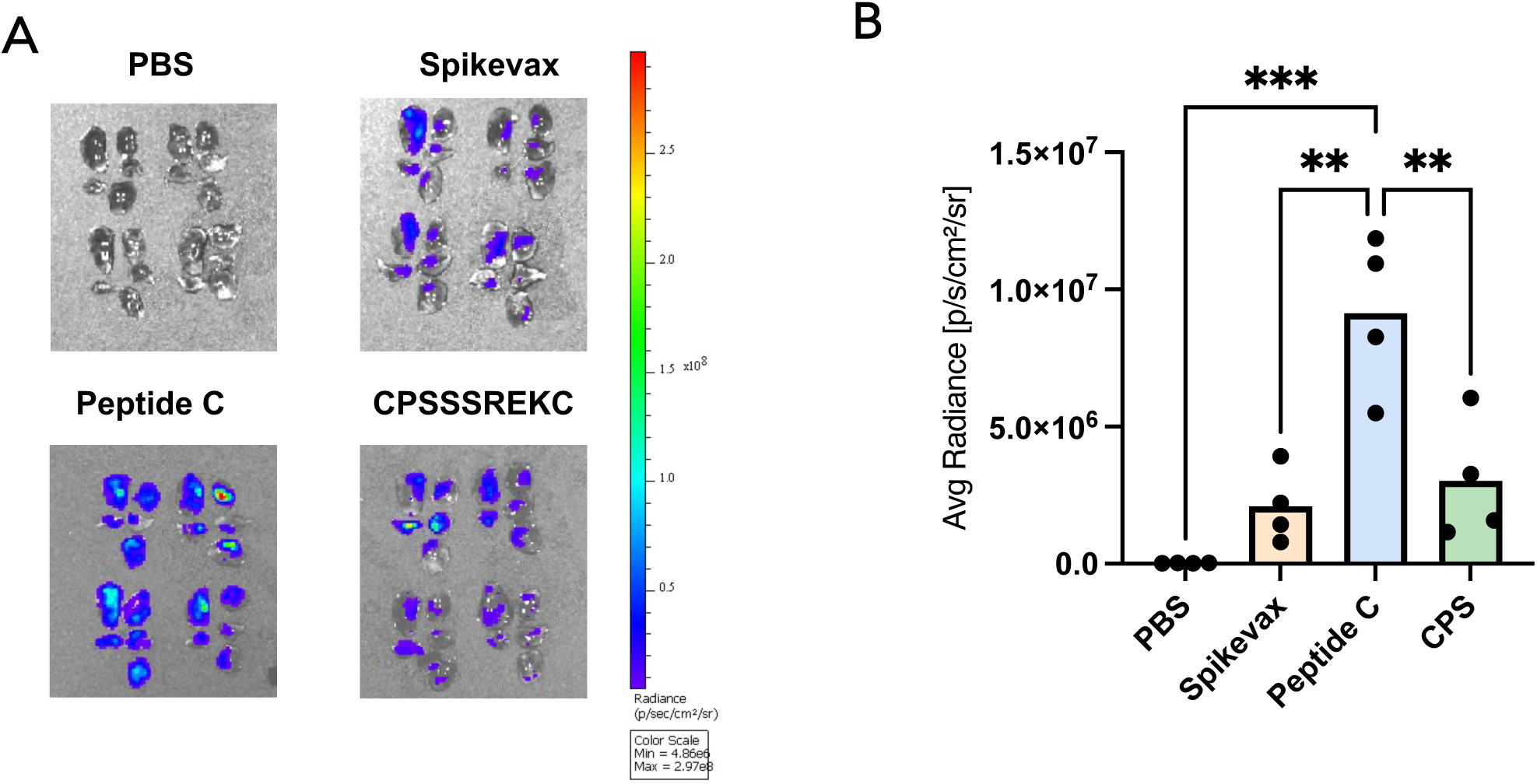
*In vivo* NLuc expression in Balb/c mice (*N*=4; 6-8 weeks old) following intratracheal administration of LNPs. **(A)** We delivered 40 μL of either PBS or LNPs formulated at 20 ng/μL, harvested lungs at 24 hours post administration, and immediately measured bioluminescence via IVIS imaging **(B)** Average Radiance (p/s/cm^2^/sr) was calculated for each lung (*N*=4) (B); One-way ANOVA, (Tukey’s multiple comparison test); *\*\*p<0.005, ***p<0.0005*.

While inclusion of our peptide is a promising first step towards targeted delivery for gene therapy of CF, other important considerations for a clinically relevant product include safety and toxicity profiles of our peptide-LNP formulation. It is important to note that since the formulations were based on the components of the COVID-19 mRNA vaccine and used a similar formulation strategy, i.e., nanoprecipitation by microfluidic mixing, our targeting approach can be easily formulated and scaled into clinical use. Additionally, given that our formulation was intratracheally instilled into mouse lungs, LNPs will need to be formulated as stable aerosols for inhalation therapy, and future studies will require optimization for aerosolization. Lastly, while we demonstrated that our peptide delivers mRNA to pHBECs, we do not have *a priori* knowledge of the target. The respiratory epithelia contain multiple cell types such as ciliated, goblet, basal, and ionocyte cells. Recent findings suggest secretory cells are thought to play a prominent role in CFTR expression, hinting therapy should be directed to this cell type. (74) While our panning allowed for unbiased identification of peptide ligands against cells, it would be important to understand which specific cell type is targeted and elucidate the mechanism of how these peptides achieve delivery. With these goals in mind, our peptide discovered through phage display and stringent selection demonstrates significant progress given that it can successfully target differentiated pHBECs and even enhance mRNA expression *in vivo*.

In this work, we implemented an improved phage display selection strategy and discovered a lead peptide that significantly increased the delivery of mRNA to fully differentiated pHBECs from CF patients and to mouse airways *in vivo*. Given that this peptide was successfully and easily implemented as a fifth component to traditional LNPs, we anticipate that it could aid in the delivery of more clinically relevant nucleic acids for CF therapy (e.g., CRISPR-Cas9 editing mRNA). Further, we conducted this work with an FDA approved LNP formulation. This demonstrates the clinical relevance of our LNP formulation. In future studies, we anticipate that our peptide could be added to other LNP systems that have been specifically optimized for pulmonary delivery. While gene therapy for CF has been a significant challenge due to poor delivery efficacy to the lungs, we are encouraged by our results presented here and see potential for our mucus and cell penetrating targeting peptide to overcome this delivery barrier.

## Materials and Methods

### Differentiation of primary human bronchial epithelial cells

For coating Transwell^®^ inserts (0.4-micron, 6.5 mm diameter, Corning Product, catalog #3470), we prepared Human Placenta Collagen Type IV (Sigma, catalog #C7521) according to protocols established by University of North Carolina, Marisco Lung Institute (MLI) Core. We first prepared a 10x stock solution (10 mg Collagen, 20 mL ddH2O, 50 μL concentrated acetic acid) and incubated from 4-8 hours at 37°C to dissolve. We then filter sterilized the solution with a 0.2 μm syringe filter, aliquotted and stored at −20°C. Prior to coating, we prepared a 1x solution with sterile cell culture grade water. Next, we added 100 μL to each insert, dried plates containing inserts (with lids off) in a biosafety cabinet overnight and UV sterilized inserts for at least 30 minutes before use.

From MLI, we purchased primary human bronchial epithelial cells (pHBECs) from seven different cystic fibrosis (CF) patients homozygous for the deltaF508 mutation (patient demographics in Supplementary Table 1). All cells used for experiments were passage 2 (P2) unless otherwise noted. We pooled all seven patients and seeded onto pre-coated inserts for differentiation at air-liquid interface (ALI) according to Pneumacult™ Ex-Plus (STEMCELL Technologies Inc., catalog #05040; supplemented with amphotericin B (0.25 μg/mL final concentration) (Thermo Fisher Scientific, catalog #BP264520), gentamicin (50 μg/mL final concentration) (Sigma, catalog #G1397), and 1x penicillin/streptomycin) and Pneumacult™ALI media (STEMCELL Technologies Inc., catalog #05001; supplemented with 1x penicillin/streptomycin) protocols. Briefly, we first expanded P2 cells on pre-coated Transwell^®^ inserts until at least 80% confluency. Once confluency was reached, we removed apical media (i.e., airlifted cells), and added Pneumacult™-ALI maintenance media to the basolateral side only. We then maintained cells in Pneumacult™-ALI maintenance media until differentiation occurred (at least 21 days post airlift and confirmed by observation with mucus production and cilia movement).

### Differentiation of THP-1 cells for macrophage uptake studies

For macrophage uptake studies, we cultured THP-1 cells (ATCC, catalog #TIB-202) according to ATCC’s recommendations. Briefly, we maintained cells in RPMI-1640 media (Sigma, catalog #R8758) supplemented with 10% Fetal Bovine Serum (FBS) and gentamicin (final concentration of 50 μg/mL). For differentiation into nonpolarized macrophages, we resuspended cells in phorbol 12-myristate 13-acetate (PMA) (Sigma, catalog #P8139) containing media (15 ng/mL) and seeded at a final density of 4E5 viable cells/mL. We seeded 0.5 mL of cell suspension per well for a 24-well plate. We incubated cells at 37°C/5 % CO_2_ for 48 hours before exchanging media with PMA-free media and incubating cells for an additional 24 hours prior to use for experiments.

### Selection Strategy

We used T7 cysteine constrained heptapeptide (CX7C) phage libraries previously made in our lab(16, 18) using T7Select415-1 cloning kit (Novagen, catalog #70015) to select against CF pHBECs to discover mucus and cell-penetrating peptides. We combined T7 libraries and diluted them in DPBS (without Ca^++^ and Mg^++^) to a final concentration of 3.3E5 plaque forming units/μL or PFU/μL to obtain 1,000 viral genomes per cell (we seeded 3.3E4 pHBECs per insert). We then added 100 μL of phage solution to the apical side of each Transwell^®^ insert *(N=*3) and incubated for a given amount of time depending on the round of selection (Round 1 = 16 hours, Round 2 = 4h, Rounds 3-5 =1h). After incubation, we removed apical and basolateral solutions, and washed cells with phage elution buffer (20 mM Tris-HCl pH 8, 100 mM NaCl, 6 mM MgSO_4_) and DPBS to remove any unbound/bound phage remaining. After wash steps, we added 200 μL of M-PER lysis buffer (Thermo Fisher Scientific Inc., catalog #78503) on top of cells and incubated for 5 minutes at 180 rotations per minute (RPM) on an orbital shaker to ensure cell lysis. Following lysis, we scraped cells (using a P200 pipette tip), collected lysate, and spun cells down for 10 min at 14,000 x g to separate cell debris from internalized phage (present in supernatant). We then quantified and titered collected phage using standard double-layer plaque assay. After each round, we pooled internalized phage from each replicate and used this pool as input for the following round. We believe this additional optimization step improved reproducibility in our selection process compared to our previous screening strategy, where we had few clones present in all three replicates.(16) Additionally, after sample collection, we reserved a portion of this sample for Next Generation Sequencing (NGS) and amplified the remaining sample in *E. coli* (BL21) until lysis was observed (∼2 hours), per manufacturer’s protocol, so input concentrations remained the same in the following rounds. As a control to identify parasitic sequences,(75) the naïve library (unselected CX7C library) was amplified 5 subsequent times prior to NGS sample preparation.

### Next Generation Sequencing Data Analysis

Following each round of selection, we isolated DNA from amplified phage, prepared samples for NGS and performed peptide sequence analysis as previously reported.(16, 18) After obtaining a frequency count of all peptide sequences from each round of panning, and in order to properly select enriched CX7C sequences from our NGS data, we omitted sequences that were linear (i.e., did not contain the expected CX7C motif) or did not contain the correct flanking sequences surrounding the insert initially cloned into our library. We then analyzed the top 30 most abundant sequences after all five rounds of selection to determine their physicochemical properties. Specifically, we calculated net charge and grand average of hydropathy score (GRAVY score) for each 7-mer sequences using *in silico* tools (https://pepcalc.com) and (http://www.gravy-calculator.de/index.php), respectively. Additionally, we created a multiple sequence alignment visual representation using Seq2Logo after we combined top 30 sequences from each replicate (total of 90 sequences) after 5 rounds of selection.(19) Peptide sequences that were found in the top 10 most frequently occurring peptide sequences, in all three replicates, and absent from an amplified naïve library control (amplified five times to correlate with the five rounds of panning) were selected for further validation studies. We evaluated the enrichment for each clone selected for validation studies by calculating the average *(N=*3) percentage of total unique sequences retrieved from NGS for each round of selection. Additionally, we analyzed peptide sequences using online tool, “Scanner and Reporter of Target-Unrelated Peptides” or SAROTUP to confirm sequences used for validation were not target-unrelated peptides.(76–78)

### Validation of Clones

After we selected peptides from NGS data for further validation, we cloned these peptide sequences back into T7 phage and verified through sanger sequencing as previously published.(18) Briefly, we obtained complementary pairs of oligonucleotides (i.e., sense and anti-sense oligos) for each peptide sequence through IDT (Supplementary Table 2). Oligos were diluted in IDTE buffer, pH 8 (IDT) to a 100 μM stock solution and then to 10 μM before use. We then annealed oligos (5 μL of 10 μM sense and anti-sense oligos in a 50 μL reaction with ultrapure water at 95°C for 3 minutes then cooled to RT (25°C) at 0.1°C/S (Ramp). After annealing, inserts were ligated to T7-Select 415-1 vector arms (Novagen (EMD Millipore), catalog #70015-3) using T4 DNA ligase (New England Biolabs, catalog #M0202L). We then packaged ligation reactions using T7Select415-1b cloning kit (Novagen (EMD Millipore), catalog #70015-3) according to manufacturer’s instructions. We plated packaged clones using a standard double-layer plaque assay with BL21 *E. coli*. After incubating clones overnight at 37°C, we isolated and sequenced individual plaques to confirm proper cloning. Once clones were validated, they were amplified in liquid BL21 *E. coli* culture prior to validation studies. To validate clones for enhanced CF pHBEC uptake, we incubated 3.3E7 PFU of each clone (in 100 μL DPBS) on the apical side of differentiated pHBECs for 1h at 37°C. Phage were collected and quantified in the same manner as our selection for round 5.

### In vitro transcription

We prepared nanoluciferase mRNA (NLuc) as previously published.(73) Briefly, we created a template for NLuc mRNA encoding sequences for a T7 promoter, 5’ UTR, codon-optimized NLuc, and 3’ UTR. We synthesized mRNA using AmpliScribe™ T7-Flash Transcription Kit (Lucigen, catalog #ASF-3507) as described previously.(79, 80) After purification with RNA Clean & Concentrator-100 (Zymo, catalog #R1019), we added a cap1 structure using the Vaccinia Capping System (NEB, M2080S) and mRNA Cap 2’-O-methyltransferase (NEB, catalog #M0366S). We then added a 3′-poly(A) tail (E. Coli Poly (A) Polymerase, NEB, catalog #M0276L). After polyadenylation, we purified mRNA again and determined mRNA concentration by Nanodrop 1000 (Thermo Fisher Scientific Inc.), and stored aliquots at −80 °C until use.

### Lipid nanoparticle synthesis

To determine if peptides discovered through phage display could be incorporated into a nanoparticle system, we used cyclic peptides conjugated to a myristic acid lipid tail (synthesized and N-terminally modified by LifeTein^®^). We used this peptide-lipid as a fifth component to Moderna’s Spikevax lipid nanoparticle formulation using a Nanoassemblr™ benchtop instrument (Precision Nanosystems Inc., Vancouver, BC, Canada). Briefly, lipids (peptide-lipid, SM-102 (Echelon, catalog #N-1102), cholesterol (Sigma, catalog #C8667), distearoylphosphatidylcholine (DSPC) (Avanti, catalog #850365), and 1,2-dimyristoyl-rac-glycero-3-methoxypolyethylene (DMG-PEG) (Avanti, catalog #880151P)) were dissolved in molecular grade ethanol at a concentration of 10 mg/mL and reporter mRNA (NLuc) was dissolved in pre-chilled sodium citrate buffer (pH 4, 100 mM). We prepared formulations at an aqueous: organic flow ratio of 3:1, flow rate of 4 mL/min, and at a volume of 500-700 μL. After formulation, LNPs were dialyzed in 1x phosphate buffered saline (PBS; pH 7.4) for 2 hours (for *in vitro* experiments) or 4 hours (for *in vivo* experiments) in 10K MWCO Slide-A-Lyzer dialysis cassettes (Thermo Fisher Scientific, catalog #87730).

### Lipid nanoparticle characterization

For characterization, we prepared LNPs in 1x PBS (pH 7.4) for size measurements (10-fold dilution) by dynamic light scattering (DLS) using Zetasizer Nano-ZS (Malvern Instruments MA, USA). To determine encapsulation efficiency percentages (EE %), we used a modified Quant-it™ RiboGreen RNA Assay Kit protocol (Thermo Fisher Scientific, Catalog # R11490). Briefly, LNPs were diluted 100-fold in 1x Tris-EDTA (TE) or 1% Triton^®^ X-100 (Thermo Fisher Scientific, catalog #BP151) to measure unencapsulated mRNA or total mRNA, respectively. We additionally prepared two low-range standard curves (one in TE and one in 1% Triton^®^). We added 100 μL of LNP sample or standard to a clear-bottom 96-well black plate (Corning, Catalog # 3631) in duplicate then proceeded to incubate all samples and standards at 37°C for 10 minutes. We then added a 2,000-fold dilution of RiboGreen Reagent and added 100 μL of the working solution to each sample or standard. After 5 minutes, we measured fluorescence using a SpectraMax M3 plate reader (Molecular Devices) at excitation/emission of 480 nm/520 nm. We calculated EE% by the following equation: [1-(unencapsulated mRNA/total mRNA)] * 100.

### Lipid nanoparticle cell uptake in vitro

To compare mRNA expression of our LNP formulations, we incubated either differentiated CF pHBECs or THP-1 derived macrophages with 450 ng of mRNA (based off EE% values) for 48 hours. We diluted LNPs in 1x PBS to a total volume of 125 μL. We measured mRNA expression by bioluminescence assay using Nano-Glo^®^ Luciferase Assay System (Promega, Catalog # N1110). Briefly, for pHBECs, we scraped cells from each treated well (remaining LNP solution left on top) and transferred to a 96-well white flat bottom plate (Corning, catalog #3912). For differentiated THP-1 cells, we removed media first, then added 50 μL of 1x PBS (without Ca^++^ or Mg^++^) to cells before scraping and collecting (to keep apical volume consistent with pHBECs). We incubated collected samples with a 1:1 ratio of Nano-Glo^®^ Luciferase Assay Buffer to lyse the cells. Nano-Glo® substrate was added to each well (one group at a time) and luminescence was read 3 minutes later using a SpectraMax M3 plate reader (Molecular Devices).

### Lipid nanoparticle delivery in vivo

We performed all procedures in accordance and under approval by the University of Texas at Austin Institutional Animal Care and Use Committee. For intratracheal administration, we first anesthetized Balb/c mice (Charles River, female, 6-8 weeks old) under 2% isoflurane before delivering 40 μL of each LNP formulation in two separate instillations of 20 μL. Following 24 hours after administration of LNPs, we euthanized mice via carbon dioxide inhalation followed by cervical dislocation and immediately harvested lungs. We briefly rinsed harvested lungs by dipping in 1x PBS and separated each lung into five separate lobes (left, cranial, middle, accessory, and caudal). We then collected lobes for each lung into a 1.5 mL microcentrifuge tube and incubated in 300 μL of Nano-Glo^®^ substrate solution for 5 minutes, before proceeding directly to imaging using an IVIS Spectrum *In Vivo* imaging system. We measured bioluminescence (average radiance [p/s/cm^2^/sr] calculated for area surrounding all five lobes) using Living Image 4.3 software (PerkinElmer).

### Statistical analysis

We performed all data analyses with GraphPad Prism 10 (GraphPad Software, La Jolla, CA) at a significance level of *p <* 0.05 unless otherwise indicated.

## Acknowledgments

We would like to especially acknowledge Leslie Fulcher, Dr. Scott Randell, and the Marisco Lung Institute/Cystic Fibrosis and Pulmonary Disease Research and Treatment Center at the University of North Carolina at Chapel-Hill for providing the primary cells used in this study.

## Funding

This work was supported by Cystic Fibrosis Foundation (GHOSH19XX0 and LEAL19XX0).

## Author contributions

**Conceptualization:** MRS, DG, JL

**Methodology**: MRS, MML, JL, RPM, SP, TD

**Investigation:** MRS, MML, YP

**Writing – Original draft:** MRS, DG

**Writing – Review & editing:** MRS, DG, MML, YP

## Competing interests

This research has a patent application by the University of Texas at Austin.

## Data and materials availability

The data for this study can be available from the corresponding author upon reasonable request.

## Notes

### Competing Interest Statement

The authors have declared no competing interest.

## References

1. Yeung JC, Machuca TN, Chaparro C, Cypel M, Stephenson AL, Solomon M, et al. Lung transplantation for cystic fibrosis. J Heart Lung Transplant. 2020;39(6):553–60.

2. McLachlan G, Alton EWFW, Boyd AC, Clarke NK, Davies JC, Gill DR, et al. Progress in Respiratory Gene Therapy. Human Gene Therapy. 2022;33(17-18):893–912.

3. Patton JS, Byron PR. Inhaling medicines: delivering drugs to the body through the lungs. Nat Rev Drug Discov. 2007;6(1):67–74.

4. Armstrong JK, Hempel G, Koling S, Chan LS, Fisher T, Meiselman HJ, Garratty G. Antibody against poly(ethylene glycol) adversely affects PEG-asparaginase therapy in acute lymphoblastic leukemia patients. Cancer. 2007;110(1):103–11.

5. Yang Q, Jacobs TM, McCallen JD, Moore DT, Huckaby JT, Edelstein JN, Lai SK. Analysis of Pre-existing IgG and IgM Antibodies against Polyethylene Glycol (PEG) in the General Population. Anal Chem. 2016;88(23):11804–12.

6. Conte G, Costabile G, Baldassi D, Rondelli V, Bassi R, Colombo D, et al. Hybrid Lipid/Polymer Nanoparticles to Tackle the Cystic Fibrosis Mucus Barrier in siRNA Delivery to the Lungs: Does PEGylation Make the Difference? ACS Appl Mater Interfaces. 2022;14(6):7565–78.

7. Ghosh D, Peng X, Leal J, Mohanty R. Peptides as drug delivery vehicles across biological barriers. J Pharm Investig. 2018;48(1):89–111.

8. Excoffon KJDA, Koerber JT, Dickey DD, Murtha M, Keshavjee S, Kaspar BK, et al. Directed evolution of adeno-associated virus to an infectious respiratory virus. Proceedings of the National Academy of Sciences. 2009;106(10):3865–70.

9. Li W, Zhang L, Johnson JS, Zhijian W, Grieger JC, Ping-Jie X, et al. Generation of novel AAV variants by directed evolution for improved CFTR delivery to human ciliated airway epithelium. Mol Ther. 2009;17(12):2067–77.

10. Sinn PL, Hwang BY, Li N, Ortiz JLS, Shirazi E, Parekh KR, et al. Novel GP64 envelope variants for improved delivery to human airway epithelial cells. Gene Ther. 2017;24(10):674–9.

11. Oyama T, Sykes KF, Samli KN, Minna JD, Johnston SA, Brown KC. Isolation of lung tumor specific peptides from a random peptide library: generation of diagnostic and cell-targeting reagents. Cancer Lett. 2003;202(2):219–30.

12. Romanczuk H, Galer CE, Zabner J, Barsomian G, Wadsworth SC, O’Riordan CR. Modification of an Adenoviral Vector with Biologically Selected Peptides: A Novel Strategy for Gene Delivery to Cells of Choice. Human Gene Therapy. 1999;10(16):2615–26.

13. Jost PJ, Harbottle RP, Knight A, Miller AD, Coutelle C, Schneider H. A novel peptide, THALWHT, for the targeting of human airway epithelia. FEBS Lett. 2001;489(2-3):263–9.

14. Writer MJ, Marshall B, Pilkington-Miksa MA, Barker SE, Jacobsen M, Kritz A, et al. Targeted gene delivery to human airway epithelial cells with synthetic vectors incorporating novel targeting peptides selected by phage display. J Drug Target. 2004;12(4):185–93.

15. Staquicini DI, Barbu EM, Zemans RL, Dray BK, Staquicini FI, Dogra P, et al. Targeted Phage Display-based Pulmonary Vaccination in Mice and Non-human Primates. Med. 2021;2(3):321–42.

16. Leal J, Peng X, Liu X, Arasappan D, Wylie DC, Schwartz SH, et al. Peptides as surface coatings of nanoparticles that penetrate human cystic fibrosis sputum and uniformly distribute in vivo following pulmonary delivery. J Control Release. 2020;322:457–69.

17. Fulcher ML, Gabriel S, Burns KA, Yankaskas JR, Randell SH. Well-differentiated human airway epithelial cell cultures. Methods Mol Med. 2005;107:183–206.

18. Mohanty RP, Liu X, Kim JY, Peng X, Bhandari S, Leal J, et al. Identification of peptide coatings that enhance diffusive transport of nanoparticles through the tumor microenvironment. Nanoscale. 2019;11(38):17664–81.

19. Thomsen MCF, Nielsen M. Seq2Logo: a method for construction and visualization of amino acid binding motifs and sequence profiles including sequence weighting, pseudo counts and two-sided representation of amino acid enrichment and depletion. Nucleic Acids Res. 2012;40(Web Server issue):W281–7.

20. Labiris NR, Dolovich MB. Pulmonary drug delivery. Part I: physiological factors affecting therapeutic effectiveness of aerosolized medications. Br J Clin Pharmacol. 2003;56(6):588–99.

21. Labiris NR, Dolovich MB. Pulmonary drug delivery. Part II: the role of inhalant delivery devices and drug formulations in therapeutic effectiveness of aerosolized medications. Br J Clin Pharmacol. 2003;56(6):600–12.

22. Williams DM. Clinical Pharmacology of Corticosteroids. Respir Care. 2018;63(6):655–70.

23. Sadiq MW, Holz O, Ellinghusen BD, Faulenbach C, Müller M, Badorrek P, et al. Lung pharmacokinetics of inhaled and systemic drugs: A clinical evaluation. Br J Pharmacol. 2021;178(22):4440–51.

24. Bhat PG, Flanagan DR, Donovan MD. Drug diffusion through cystic fibrotic mucus: steady-state permeation, rheologic properties, and glycoprotein morphology. J Pharm Sci. 1996;85(6):624–30.

25. Wine JJ. The genesis of cystic fibrosis lung disease. J Clin Invest. 1999;103(3):309–12.

26. Sanders NN, De Smedt SC, Van Rompaey E, Simoens P, De Baets F, Demeester J. Cystic Fibrosis Sputum. Am J Respir Crit Care Med. 2000;162(5):1905–11.

27. Dawson M, Wirtz D, Hanes J. Enhanced viscoelasticity of human cystic fibrotic sputum correlates with increasing microheterogeneity in particle transport. J Biol Chem. 2003;278(50):50393–401.

28. Suk JS, Lai SK, Wang Y-Y, Ensign LM, Zeitlin PL, Boyle MP, Hanes J. The penetration of fresh undiluted sputum expectorated by cystic fibrosis patients by non-adhesive polymer nanoparticles. Biomaterials. 2009;30(13):2591–7.

29. Allaire NE, Griesenbach U, Kerem B, Lueck JD, Stanleigh N, Oren YS. Gene, RNA, and ASO-based therapeutic approaches in Cystic Fibrosis. J Cyst Fibros. 2023;22 Suppl 1(Suppl 1):S39–S44.

30. Bosquillon C, Madlova M, Patel N, Clear N, Forbes B. A Comparison of Drug Transport in Pulmonary Absorption Models: Isolated Perfused rat Lungs, Respiratory Epithelial Cell Lines and Primary Cell Culture. Pharm Res. 2017;34(12):2532–40.

31. Rayner RE, Makena P, Prasad GL, Cormet-Boyaka E. Optimization of Normal Human Bronchial Epithelial (NHBE) Cell 3D Cultures for in vitro Lung Model Studies. Sci Rep. 2019;9(1):500.

32. Rayner RE, Wellmerling J, Osman W, Honesty S, Alfaro M, Peeples ME, Cormet-Boyaka E. In vitro 3D culture lung model from expanded primary cystic fibrosis human airway cells. J Cyst Fibros. 2020;19(5):752–61.

33. Scudieri P, Musante I, Venturini A, Guidone D, Genovese M, Cresta F, et al. Ionocytes and CFTR Chloride Channel Expression in Normal and Cystic Fibrosis Nasal and Bronchial Epithelial Cells. Cells. 2020;9(9).

34. Cidem A, Bradbury P, Traini D, Ong HX. Modifying and Integrating in vitro and ex vivo Respiratory Models for Inhalation Drug Screening. Front Bioeng Biotechnol. 2020;8:581995.

35. Silva IAL, Laselva O, Lopes-Pacheco M. Advances in Preclinical In Vitro Models for the Translation of Precision Medicine for Cystic Fibrosis. J Pers Med. 2022;12(8).

36. Stewart CE, Torr EE, Mohd Jamili NH, Bosquillon C, Sayers I. Evaluation of differentiated human bronchial epithelial cell culture systems for asthma research. J Allergy (Cairo). 2012;2012:943982.

37. Herman RE, Makienko EG, Prieve MG, Fuller M, Houston ME, Johnson PH. Phage display screening of epithelial cell monolayers treated with EGTA: identification of peptide FDFWITP that modulates tight junction activity. J Biomol Screen. 2007;12(8):1092–101.

38. Selection and Screening Strategies. In: Geyer SSSCR, editor. Phage Display In Biotechnology and Drug Discovery. 2 ed: CRC Press; 2015. p. 97–111.

39. Barry MA, Dower WJ, Johnston SA. Toward cell-targeting gene therapy vectors: selection of cell-binding peptides from random peptide-presenting phage libraries. Nat Med. 1996;2(3):299–305.

40. Kumari A, Pal S, G BR, Mohny FP, Gupta N, Miglani C, et al. Surface-Engineered Mucus Penetrating Nucleic Acid Delivery Systems with Cell Penetrating Peptides for the Lungs. Mol Pharm. 2022;19(5):1309–24.

41. Schuster BS, Suk JS, Woodworth GF, Hanes J. Nanoparticle diffusion in respiratory mucus from humans without lung disease. Biomaterials. 2013;34(13):3439–46.

42. Xu Q, Ensign LM, Boylan NJ, Schön A, Gong X, Yang J-C, et al. Impact of Surface Polyethylene Glycol (PEG) Density on Biodegradable Nanoparticle Transport in Mucus ex Vivo and Distribution in Vivo. ACS Nano. 2015;9(9):9217–27.

43. Maisel K, Reddy M, Xu Q, Chattopadhyay S, Cone R, Ensign LM, Hanes J. Nanoparticles coated with high molecular weight PEG penetrate mucus and provide uniform vaginal and colorectal distribution in vivo. Nanomedicine. 2016;11(11):1337–43.

44. Li H, Islem Guissi NE, Su Z, Ping Q, Sun M. Effects of surface hydrophilic properties of PEG-based mucus-penetrating nanostructured lipid carriers on oral drug delivery. RSC Advances. 2016;6(87):84164–76.

45. Schneider CS, Xu Q, Boylan NJ, Chisholm J, Tang BC, Schuster BS, et al. Nanoparticles that do not adhere to mucus provide uniform and long-lasting drug delivery to airways following inhalation. Science Advances. 2017;3(4):e1601556.

46. Li P, Chen X, Shen Y, Li H, Zou Y, Yuan G, et al. Mucus penetration enhanced lipid polymer nanoparticles improve the eradication rate of Helicobacter pylori biofilm. Journal of Controlled Release. 2019;300:52–63.

47. Qiu Y, Man RCH, Liao Q, Kung KLK, Chow MYT, Lam JKW. Effective mRNA pulmonary delivery by dry powder formulation of PEGylated synthetic KL4 peptide. Journal of Controlled Release. 2019;314:102–15.

48. Futaki S, Suzuki T, Ohashi W, Yagami T, Tanaka S, Ueda K, Sugiura Y. Arginine-rich Peptides: an abundant source of membrane-permeable peptides having potential as carriers for intracellular protein delivery. Journal of Biological Chemistry. 2001;276(8):5836–40.

49. Milletti F. Cell-penetrating peptides: classes, origin, and current landscape. Drug Discovery Today. 2012;17(15):850–60.

50. Li LD, Crouzier T, Sarkar A, Dunphy L, Han J, Ribbeck K. Spatial Configuration and Composition of Charge Modulates Transport into a Mucin Hydrogel Barrier. Biophysical Journal. 2013;105(6):1357–65.

51. Marczynski M, Käsdorf BT, Altaner B, Wenzler A, Gerland U, Lieleg O. Transient binding promotes molecule penetration into mucin hydrogels by enhancing molecular partitioning. Biomater Sci. 2018;6(12):3373–87.

52. Cheng Q, Wei T, Farbiak L, Johnson LT, Dilliard SA, Siegwart DJ. Selective organ targeting (SORT) nanoparticles for tissue-specific mRNA delivery and CRISPR– Cas gene editing. Nature Nanotechnology. 2020;15(4):313–20.

53. Kwon EJ, Skalak M, Lo Bu R, Bhatia SN. Neuron-Targeted Nanoparticle for siRNA Delivery to Traumatic Brain Injuries. ACS Nano. 2016;10(8):7926–33.

54. Lo JH, Hao L, Muzumdar MD, Raghavan S, Kwon EJ, Pulver EM, et al. iRGD-guided Tumor-penetrating Nanocomplexes for Therapeutic siRNA Delivery to Pancreatic Cancer. Molecular Cancer Therapeutics. 2018;17(11):2377–88.

55. Jain PK, Lo JH, Rananaware S, Downing M, Panda A, Tai M, et al. Non-viral delivery of CRISPR/Cas9 complex using CRISPR-GPS nanocomplexes. Nanoscale. 2019;11(44):21317–23.

56. Munir M, Kett VL, Dunne NJ, McCarthy HO. Development of a Spray-Dried Formulation of Peptide-DNA Nanoparticles into a Dry Powder for Pulmonary Delivery Using Factorial Design. Pharm Res. 2022;39(6):1215–32.

57. Herrera-Barrera M, Ryals RC, Gautam M, Jozic A, Landry M, Korzun T, et al. Peptide-guided lipid nanoparticles deliver mRNA to the neural retina of rodents and nonhuman primates. Science Advances. 2023;9(2):eadd4623.

58. Anthiya S, Öztürk SC, Yanik H, Tavukcuoglu E, Şahin A, Datta D, et al. Targeted siRNA lipid nanoparticles for the treatment of KRAS-mutant tumors. Journal of Controlled Release. 2023;357:67–83.

59. Roces CB, Lou G, Jain N, Abraham S, Thomas A, Halbert GW, Perrie Y. Manufacturing Considerations for the Development of Lipid Nanoparticles Using Microfluidics. Pharmaceutics. 2020;12(11):1095.

60. Hisert KB, Liles WC, Manicone AM. A Flow Cytometric Method for Isolating Cystic Fibrosis Airway Macrophages from Expectorated Sputum. Am J Respir Cell Mol Biol. 2019;61(1):42–50.

61. Hou F, Xiao K, Tang L, Xie L. Diversity of Macrophages in Lung Homeostasis and Diseases. Front Immunol. 2021;12:753940.

62. Wright AKA, Rao S, Range S, Eder C, Hofer TPJ, Frankenberger M, et al. Pivotal Advance: Expansion of small sputum macrophages in CF: failure to express MARCO and mannose receptors. J Leukoc Biol. 2009;86(3):479–89.

63. Murphy BS, Bush HM, Sundareshan V, Davis C, Hagadone J, Cory TJ, et al. Characterization of macrophage activation states in patients with cystic fibrosis. J Cyst Fibros. 2010;9(5):314–22.

64. Ruge CA, Kirch J, Cañadas O, Schneider M, Perez-Gil J, Schaefer UF, et al. Uptake of nanoparticles by alveolar macrophages is triggered by surfactant protein A. Nanomedicine: Nanotechnology, Biology and Medicine. 2011;7(6):690–3.

65. Ruge CA, Schaefer UF, Herrmann J, Kirch J, Cañadas O, Echaide M, et al. The Interplay of Lung Surfactant Proteins and Lipids Assimilates the Macrophage Clearance of Nanoparticles. PLOS ONE. 2012;7(7):e40775.

66. Geiser M, Quaile O, Wenk A, Wigge C, Eigeldinger-Berthou S, Hirn S, et al. Cellular uptake and localization of inhaled gold nanoparticles in lungs of mice with chronic obstructive pulmonary disease. Particle and Fibre Toxicology. 2013;10(1):19.

67. Yin B, Chan CKW, Liu S, Hong H, Wong SHD, Lee LKC, et al. Intrapulmonary Cellular-Level Distribution of Inhaled Nanoparticles with Defined Functional Groups and Its Correlations with Protein Corona and Inflammatory Response. ACS Nano. 2019;13(12):14048–69.

68. Lokugamage MP, Vanover D, Beyersdorf J, Hatit MZC, Rotolo L, Echeverri ES, et al. Optimization of lipid nanoparticles for the delivery of nebulized therapeutic mRNA to the lungs. Nat Biomed Eng. 2021;5(9):1059–68.

69. Shimosakai R, Khalil IA, Kimura S, Harashima H. mRNA-Loaded Lipid Nanoparticles Targeting Immune Cells in the Spleen for Use as Cancer Vaccines. Pharmaceuticals (Basel). 2022;15(8):1017.

70. Leppek K, Byeon GW, Kladwang W, Wayment-Steele HK, Kerr CH, Xu AF, et al. Combinatorial optimization of mRNA structure, stability, and translation for RNA-based therapeutics. Nat Commun. 2022;13(1):1536.

71. Kimura S, Harashima H. On the mechanism of tissue-selective gene delivery by lipid nanoparticles. Journal of Controlled Release. 2023.

72. Huayamares SG, Lokugamage MP, Rab R, Da Silva Sanchez AJ, Kim H, Radmand A, et al. High-throughput screens identify a lipid nanoparticle that preferentially delivers mRNA to human tumors in vivo. Journal of Controlled Release. 2023;357:394–403.

73. Lewis MM, Soto MR, Maier E, Wulfe S, Bakheet S, Obregon H, Ghosh D. Optimization of ionizable lipids for aerosolizable mRNA lipid nanoparticles. Bioengineering & Translational Medicine. 2023;8(5).

74. Okuda K, Dang H, Kobayashi Y, Carraro G, Nakano S, Chen G, et al. Secretory Cells Dominate Airway CFTR Expression and Function in Human Airway Superficial Epithelia. Am J Respir Crit Care Med. 2021;203(10):1275–89.

75. Liu GW, Livesay BR, Kacherovsky NA, Cieslewicz M, Lutz E, Waalkes A, et al. Efficient Identification of Murine M2 Macrophage Peptide Targeting Ligands by Phage Display and Next-Generation Sequencing. Bioconjug Chem. 2015;26(8):1811–7.

76. Huang J, Ru B, Li S, Lin H, Guo F-B. SAROTUP: scanner and reporter of target-unrelated peptides. J Biomed Biotechnol. 2010;2010:101932.

77. Huang J, Ru B, Zhu P, Nie F, Yang J, Wang X, et al. MimoDB 2.0: a mimotope database and beyond. Nucleic Acids Res. 2012;40(Database issue):D271–7.

78. He B, Chai G, Duan Y, Yan Z, Qiu L, Zhang H, et al. BDB: biopanning data bank. Nucleic Acids Res. 2016;44(D1):D1127–32.

79. Kauffman KJ, Dorkin JR, Yang JH, Heartlein MW, DeRosa F, Mir FF, et al. Optimization of Lipid Nanoparticle Formulations for mRNA Delivery in Vivo with Fractional Factorial and Definitive Screening Designs. Nano Lett. 2015;15(11):7300–6.

80. Zeng C, Hou X, Yan J, Zhang C, Li W, Zhao W, et al. Leveraging mRNA Sequences and Nanoparticles to Deliver SARS-CoV-2 Antigens In Vivo. Adv Mater. 2020;32(40):e2004452.

