## Supplemental Information for "Discovery of peptides for targeted delivery of mRNA lipid nanoparticles to cystic fibrosis lung epithelia"

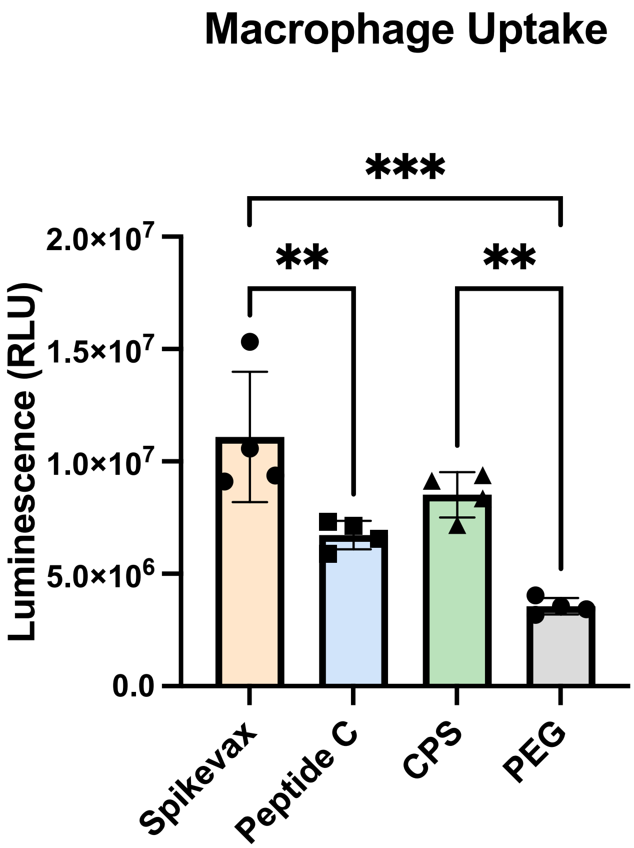

**Fig. S1.**

**NLuc mRNA expression in THP-1 derived macrophages following LNP treatment.** We added LNPs (450 ng of mRNA) to THP-1 derived macrophages (*N*=4 per treatment group) and incubated them at 37°C for 48 hours. Following incubation, we measured bioluminescence by plate reader. One-way ANOVA (Tukey's multiple comparisons test; *** p <0.01, *** p<0.005.*

**Table S1.**

| Patient | Age | Sex | Race |
|---|---|---|---|
| 1 | 24 | F | Unknown |
| 2 | 28 | M | Unknown |
| 3 | 46 | F | Caucasian |
| 4 | 20 | M | Caucasian |
| 5 | 29 | F | Unknown |
| 6 | 30 | M | Unknown |
| 7 | 38 | F | Unknown |

**List of patient demographics.** All patients are homozygous for deltaF508 mutation.

**Table S2.**

| Clone Identifier | Sequences | Oligonucleotides |
|---|---|---|
| A | CTPKRSRAC | Sense: 5' --- AAT TCT TGC ACA CCC AAA AGG AGT AGA GCT TGC TAA A --- 3' |
| | | Asn: 5' --- AGCTT TTA GCA AGC TCT ACT CCT TTT GGG TGT GCA AG --- 3' |
| B | CTRPTRSKC | Sense: 5' --- AAT TCT TGC ACA AGG CCC ACT AGA AGT AAA TGC TAA A --- 3' |
| | | Asn: 5' --- AGCTT TTA GCA TTT ACT TCT AGT GGG CCT TGT GCA AG --- 3' |
| C | CTSTRKKQC | Sense: 5' --- AAT TCT TGC ACT TCA ACA AGG AAG AAA CAA TGC TAA A --- 3' |
| | | Asn: 5' --- AGCTT TTA GCA TTG TTT CTT CCT TGT TGA AGT GCA AG--- 3' |
| D | CPAPRGKRC | Sense: 5' --- AAT TCT TGC CCC GCT CCA AGG GGA AAA AGA TGC TAA A --- 3' |
| | | Asn: 5' --- AGCTT TTA GCA TCT TTT TCC CCT TGG AGC GGG GCA AG --- 3' |
| E | CAPSKRNRC | Sense: 5' --- AAT TCT TGC GCT CCC AGT AAA AGA AAT AGG TGC TAA A --- 3' |
| | | Asn: 5' --- AGCTT TTA GCA CCT ATT TCT TTT ACT GGG AGC GCA AG --- 3' |
| F | CLSPTGKAC | Sense: 5' --- AAT TCT TGC CTA AGT CCC ACA GGA AAA GCT TGC TAA A --- 3' |
| | | Asn: 5' --- AGCTT TTA GCA AGC TTT TCC TGT GGG ACT TAG GCA AG --- 3' |

**Oligonucleotide Sequences used for validation studies.** Sequences were optimized using GenSmart Codon Optimization Tool. Restriction Site (to be joined with 415-b vector arms that are precut); Inserted to make insert in reading frame; Stop Codon; Cysteine. Note: Clone CPS was obtained from a previously made stock in our lab.(*16*) Asn = Anti-sense.
